# Enhancement of response learning in male rats with intrastriatal infusions of a BDNF - TrkB agonist, 7,8-dihydroxyflavone

**DOI:** 10.1101/2024.08.08.606692

**Authors:** Robert S. Gardner, Matthew T. Ambalavanar, Paul E. Gold, Donna L. Korol

## Abstract

Enhancement of learning and memory by cognitive and physical exercise may be mediated by brain-derived neurotrophic factor (BDNF) acting at tropomyosin receptor kinase B (TrkB). Upregulation of BDNF and systemic administration of a TrkB agonist, 7,8-dihydroxyflavone (7,8-DHF), enhance learning of several hippocampus-sensitive tasks in rodents. Although BDNF and 7,8-DHF enhance functions of other brain areas too, these effects have mainly targeted non-cognitive functions. One goal of the present study was to determine whether 7,8-DHF would act beyond the hippocampus to enhance cognitive functions sensitive to manipulations of the striatum. Here, we examined the effects of intrastriatal infusions of 7,8-DHF on learning a striatum-sensitive response maze and on phosphorylation of TrkB receptors in 3-month-old male Sprague Dawley rats. Most prior studies of BDNF and 7,8-DHF effects on learning and memory have administered the drugs for days to months before assessing effects on cognition. A second goal of the present study was to determine whether a single drug treatment near the time of training would effectively enhance learning. Moreover, 7,8-DHF is often tested for its ability to reverse impairments in learning and memory rather than to enhance these functions in the absence of impairments. Thus, a third goal of this experiment was to evaluate the efficacy of 7,8-DHF in enhancing learning in unimpaired rats. In untrained rats, intrastriatal infusions of 7,8-DHF resulted in phosphorylation of TrkB receptors, suggesting that 7,8-DHF acted as a TrkB agonist and BDNF mimic. The findings that a single, intra-striatal infusion of 7,8-DHF 20 min before training enhanced response learning in rats suggest that, in addition to its trophic effects, BDNF modulates learning and memory through receptor mediated cell signaling events.

## 1. Introduction

Brain-derived neurotrophic factor (BDNF) is known for its role as a regulator of learning, memory, and synaptic plasticity (e.g., Gonzales et al., 2019; Leal et al., 2017; Lu et al., 2014; Miranda et al., 2019). BDNF enhancement of memory appears to be mediated primarily by actions at the BDNF receptor, tropomyosin receptor kinase B (TrkB) (for reviews see: Bekinschtein et al., 2014; Cunha et al., 2010; Numakawa and Odaka, 2022; Vivar et al., 2013; Yang and Zhu, 2022). Of note, BDNF levels and TrkB phosphorylation in the hippocampus are positively correlated with acquisition of spatial learning and memory tasks, while genetic manipulations or pharmacological inhibition of BDNF and TrkB signaling lead to learning and memory deficits (Minichiello et al., 2002; Mizuno et al., 2000, 2003; Pang et al., 2024; Tyler et al., 2002). Other interventions such as environment enrichment, voluntary physical exercise, and priming with prior cognitive training, each increase brain BDNF levels and enhance learning and memory in a BDNF or TrkB-dependent manner (Korol et al., 2013; Novkovic et al., 2015; van Praag et al., 1999; Vaynman et al., 2004; Xu et al., 2021).

Of interest here, 7,8-dihydroxyflavone (7,8-DHF) is an isoflavone that, when injected or ingested, activates TrkB receptors thereby mimicking some of the effects of BDNF (Jang et al., 2010; Liu et al., 2014; Yang and Zhu, 2022). Systemic administration of 7,8-DHF enhances learning, as tested mainly in hippocampus-sensitive (e.g., Agrawal et al, 2015; Bollen et al., 2013; Castello et al., 2014; Giacomini et al., 2017; Huang et al., 2021; Pandey et al., 2020; Seese et al., 2020; Zhang et al., 2022) and amygdala-sensitive (Andero et al., 2011; Ou and Gean, 2006; Rattiner et al., 2004; Zeng et al., 2012a) tasks.

One goal of the present study was to determine whether 7,8-DHF would also enhance learning a striatum-sensitive task, specifically when the drug was administered directly to the striatum. The dorsolateral striatum is engaged when individuals learn tasks using habit, egocentric, or response strategies (Gahnstrom and Spiers, 2020; Gold, 2016; Gold et al., 2013; Gomez-Perales and Brake, 2023; Goodman and McIntyre, 2017; Graybiel, 2016; Iaria et al., 2003; Korol, 2004; Korol et al., 2013; Mizumori and Jo, 2013; Packard and Goodman, 2013; Packard and McGaugh, 1996; Packard et al., 2021; Sodums and Bohbot, 2020; White and Bohbot, 2017; White and McDonald, 2002; White et al., 2013). Although, to our knowledge, there are no prior studies of BDNF contributions to striatal functions within a memory system framework, there is evidence that BDNF acting through TrkB receptors reverses non-mnemonic striatal dysfunctions. For example, daily systemic administration of 7,8-DHF delays or ameliorates motor deficits in rodent models of Parkinson’s and Huntington’s diseases (Garcia-Diaz Barriga et al., 2017; Lee and Han, 2019; Li et al., 2016; Massaquoi et al., 2020; Sconce et al. 2015; Zuo et al., 2021).

Exercise is a well-documented intervention that enhances learning and memory in many tasks (for reviews see: Bettio et al., 2019; Erickson et al., 2019; Jaberi and Fahnestock, 2023; Rendeiro and Rhodes, 2018). We have found that prior physical or cognitive activity enhanced learning of both hippocampus-and striatum-sensitive tasks (Korol et al., 2013). Specifically, voluntary wheel running or performing a working memory task accelerated subsequent learning of mazes designed to be solved readily using either place or response solutions, i.e., solutions based on hippocampal or striatal strategies, respectively. In addition to enhanced learning, these priming experiences resulted in increased BDNF concentrations in both the hippocampus and striatum. Particularly relevant to the current study, pretraining infusions of inhibitors of BDNF signaling into the hippocampus and striatum blocked the enhancing effects of the priming experiences on the canonical place and response learning tasks, respectively (Korol et al., 2013; Vaynman et al., 2004). Together with the reported correlation between phosphorylated TrkB in the dorsal striatum and response learning efficacy (Pahng and Colombo, 2017), the findings suggest that BDNF activation of TrkB receptors in the striatum at the time of testing contributes to striatum-sensitive learning.

Here we tested in male rats the effects of activation of the TrkB receptor with intrastriatal infusions of 7,8-DHF on learning a response maze task we have used often to test striatal contributions to learning and memory (Chang and Gold, 2004; Gardner et al., 2020; Gold et al., 2013; Korol and Pisani, 2015; Pych et al., 2005; Scavuzzo et al., 2021; Zurkovsky et al., 2011).

A second goal of the present study was to test the acute effects of 7,8-DHF when the drug is administered once near the time of training and thus without its long-term trophic effects. Most previous studies of 7,8-DHF have used chronic treatments, e.g., daily injections for several days to weeks before behavioral testing (e.g., Agrawal et al., 2015; Akhtar et al., 2021; Bryant et al., 2021; Castello et al., 2014; Krishna et al., 2017; Seese et al., 2020; Stagni et al., 2017). The use of extended treatment durations was in part to reverse learning and memory deficits in long-term models of dysfunction, e.g. rodent models of traumatic brain injury (Agrawal et al., 2015), Fragile X syndrome (Seese et al., 2020; Tian et al., 2015), Down syndrome (Giacomini et al., 2019; Stagni et al., 2017), aging (Zeng et al., 2012b), or Alzheimer’s disease (Chen et al., 2014; Devi and Ohno, 2012; Gao et al., 2016; Zhang et al., 2014). We found few reports showing enhancement of learning with a single administration of 7,8-DHF near the time of testing (e.g., Andero et al., 2011; Bollen et al., 2013). Here, the drug was administered once by direct infusion into the striatum 20 min before training.

In addition to determining whether a single infusion of 7,8-DHF would rapidly enhance learning, a third goal was to determine whether the drug can enhance learning and memory in rats without a cognitive deficit. Although 7,8-DHF can act on striatal processes to augment non-cognitive functions to delay or to reduce motor deficits in rodent models of Parkinson’s and Huntington’s diseases, drug enhancement of motor functions was not evident in control groups (Garcia-Diaz Barriga et al., 2017; Li et al., 2016; Massaquoi et al., 2020; Sconce et al. 2015). The absence of effects in control groups opens the possibility that BDNF regulation of striatum-based cognition will also be specific to contexts where deficits are expected.

To complement the effects of intrastriatal infusions of 7,8-DHF on learning, we verified 7,8-DHF-induced activation of striatal TrkB receptors by measuring phosphorylated tyrosine 816 after the drug infusions. Phosphorylation of tyrosine 816 on TrkB receptors (pTrkB^Y816^) is associated with BDNF and 7,8-DHF signaling through TrkB with downstream CREB activation and is implicated in TrkB-dependent effects on learning and memory (Minichiello, 2009; Zeng et al., 2012a, b). We found that acute, intracerebral treatments of 7,8-DHF improved response learning and increased TrkB phosphorylation.

## 2. Methods

### 2.1 Animals

Three-month-old male Sprague Dawley rats obtained from Envigo (Indianapolis, IN, United States) were used for these experiments. Upon arrival, rats were individually housed in a room with a 12 hr light/12 hr dark cycle for at least two days prior to any experimental procedures. All methods were carried out in accordance with the National Institutes of Health Guide for the Care and Use of Laboratory Animals and approved by the Syracuse University Institutional Animal Care and Use Committee.

### 2.2 Experimental design

Briefly, rats underwent cranial surgery followed by food restriction after a week of post-operative recovery. One set of rats received drug infusions into the dorsolateral striatum 20 min prior to response training. After training, these rats were euthanized, and their brains processed for histology. A second set of rats that did not undergo training received drug infusions 20 min prior to tissue collection. In these rats, the dorsal striata from both hemispheres were processed for western blot analyses of TrkB activation.

### 2.3 Cranial Surgery

Rats were anesthetized with isoflurane (2-4%) and placed in a stereotaxic apparatus with a nosepiece that provided isoflurane delivery during surgery using a SomnoSuite anesthesia system (Kent Scientific, Torrington, CT). Stainless steel guide cannulae (22-gauge; Plastics One, Roanoke VA) were implanted bilaterally into the dorsolateral striatum (coordinates: 0.2 mm anterior and 3.6 mm lateral to bregma, 2.8 mm ventral to dura). Dental cement and head screws affixed the cannulae to the skull. Pre-operative care included analgesic (Flunixin: 2.5 mg / kg; subcutaneous) and antibiotic (Penicillin: 100,000 units / kg; intramuscular). Immediately after surgery, rats received an injection of saline (10 ml; subcutaneous). In addition, ibuprofen (47 mg / 500 ml) was added to the rats’ water bottles for several days after surgery. Health and well-being were monitored daily for seven days after surgery; no ill postoperative effects were evident.

### 2.4 Food Restriction

Beginning at least one week after surgery, rats were handled daily (3-5 min) and food-restricted (7-10 grams of rat chow provided each day) to reach 80-83% of their free-feeding weights over the course of 7-9 days. On each day of food restriction, rats were given three Frosted Cheerios® (General Mills), the food reward using during response training.

### 2.5 7,8-DHF and vehicle infusions

On the day of training or tissue collection, stainless steel infusion cannulae (28-gauge; Plastics One), extending 1 mm past the guide cannulae, were inserted bilaterally. Either 7,8-DHF or vehicle (78.7% DMSO in saline) was infused bilaterally into the dorsolateral striatum (0.5 µl / hemisphere) with a micro-syringe pump at a rate of 0.25 µl/min. Rats that underwent training received bilateral infusions of vehicle (n = 4) or one of three doses of 7,8-DHF (Sigma-Aldrich, St. Louis, MO): 1 µg (n = 4), 5 µg (n = 4), or 10 µg (n = 4) per hemisphere. Infusion needles were left in place for 1 min after infusions to allow diffusion of 7,8-DHF away from the cannulae. After the infusions, rats were placed into a clean transfer cage and acclimated to the training room for 20 min before testing. For western blot experiments, two additional groups of untrained rats received intrastriatal infusions as above with either 1 µg 7,8-DHF (n = 8) or vehicle (n = 8).

### 2.6 Response training

Training began 20 min after the end of the infusions. Rats were trained in a single session (75 trials; 30-sec inter-trial interval) to find food (1/2 Frosted Cheerios^®^) on a maze using a response solution, i.e., a turning-based rule as described previously (Chang and Gold, 2004; Gardner et al., 2020; Korol and Kolo, 2002; Matthews and Best, 1995; Newman et al., 2017; Packard and McGaugh, 1996; Scavuzzo et al., 2021). The training apparatus was a plus-shaped 4-arm maze configured into a T by obstructing one arm with a removable blockade (Figure 1). Each arm of the maze was 45 cm long, 14 cm wide, and had 7.5 cm tall walls; a center choice area measured 14 x 14 cm. Reward cups were placed at the ends of each arm. The maze was surrounded by a solid curtain to minimize extra-maze visual cues. On each trial, the rat started from one of two possible positions and was rewarded for making a consistent turn (e.g., turning right; Figure 1). To limit the use of alternate strategies (e.g., spatial or olfactory), several steps were taken: 1) the start location was pseudo-randomly assigned and counterbalanced across the 75-trial training session with the rule that no more than three consecutive trials were started from the same arm, 2) each reward cup contained inaccessible reward, and 3) the maze was rotated 90 degrees between each trial.

**Figure 1.**
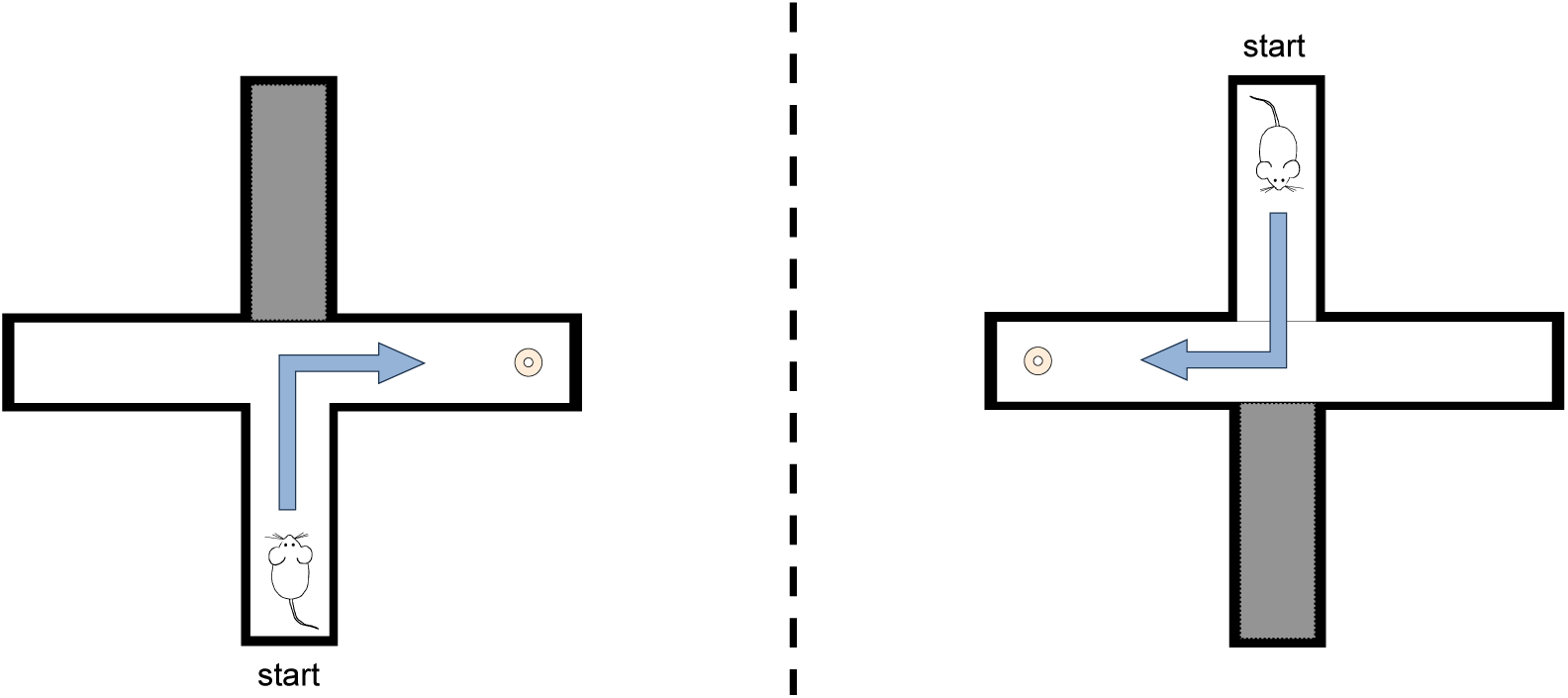
Illustration of maze use in response training. Response training was conducted in a single session (75-trials; 30 sec inter-trial-interval) on a T-maze. Rats received a reward for making a consistent body turn from one of two distinct starting positions. The start positions were quasi-randomly assigned across trials and counterbalanced across the session. Arrows indicate the rewarded turn for each starting position.

To start a trial, the experimenter placed the rat on the assigned start arm for each trial and remained standing behind that arm during the trial. The trial was completed when the rat made a choice and traveled to the end of the chosen arm or after two min, whichever came first. A choice was recorded when all four paws crossed from the center choice area into the arm. When the rat chose the rewarded arm, it was allowed to consume the reward before being removed from the maze and placed in a holding cage with bedding and water during the 30-sec intertrial interval. If the rat chose the incorrect arm, it was allowed to remain at the end of the arm for 10 sec or until it turned to leave before being removed from the maze and placed in the holding cage. Between trials, the maze was rotated and replenished with food reward if necessary.

### 2.7 Tissue collection

For cannula placement verification, rats in the first set were given an overdose of sodium pentobarbital (50 mg/kg, i.p.; Vortech Pharmaceuticals, Dearborn, MI) immediately after training and then perfused with 0.9% saline followed by 10% formalin. Brains were removed and post-fixed in 10% formalin for 1-2 days at 4°C. Brains were subsequently transferred to 20% glycerol in phosphate-buffered saline before being sectioned (30 µM; coronal plane) and stained with cresyl violet. Injection sites were confirmed with light microscopy.

Rats in the second set received an overdose of sodium pentobarbital (50 mg/rat) twenty min after infusions of either vehicle or 1 µg / hemisphere 7,8-DHF infusions (n = 8 / group). Rats were decapitated, brains were removed, and the dorsal striata were quickly dissected and frozen on dry ice. Samples were subsequently pulverized on dry ice using a mortar and pestle, aliquoted, and stored at −80°C until preparation for western blot analyses of total TrkB and phosphorylated TrkB^Y816^ proteins.

### 2.8 Western blotting

Aliquots of striata were suspended in homogenization buffer (1 mg tissue: 3 µl RIPA buffer; Santa Cruz: 24948) and homogenized (Polytron PT 1600 E; max speed) on ice. Samples were centrifuged (10,000 x g; 5 min at 4°C) and the supernatant collected. To maximize membrane protein extraction, the pellet was re-suspended in RIPA buffer (half of the original volume used for homogenization), vortexed every 5 min over a 30 min duration, and centrifuged, followed by collection of supernatant. This process was repeated once more (using one quarter of the RIPA volume used for homogenization). These supernatants were combined to form each individual experimental sample, total protein of which was measured using the Pierce BCA assay. Subsequently, the sample was mixed 1:4 with Protein Loading Buffer (4.66 ml glycerol, 1.33 ml dH_2_O, 9.33 mg BPS, 2.67ml 10% SDS, 1.33 ml β-mercaptoethanol) and heated to 99°C degrees for 7 min. An aliquot from each individual sample was combined to form a pooled sample. Pooled samples were used to generate curves to standardize readings across blots.

A concentration of 3 µg / µl total protein was used for western blot experiments. Each gel included individual samples from both treatment conditions, All Blue Prestained Protein Standards (Bio-Rad), and four pooled samples of increasing protein amounts for a standard curve (10, 20, 30, and 40 µg). Each experimental sample was evaluated in duplicate across two separate western blotting runs. Samples (30 µg total protein) were loaded onto Criterion TGX 4-20% gels (Bio-Rad: 5671094) and resolved via electrophoresis (90 V for 2 hr). Protein was subsequently transferred to methanol-activated PVDF membranes (400 mA for 45 min). Successful transfer was assessed using Ponceau total protein stain (1 min incubation) after which membranes were washed in dH_2_O followed by tris-buffered saline (TBS).

Membranes were blocked for 1 hr in TBS Odyssey blocking buffer (LI-COR) and incubated with anti-phospho-TrkB ^Y816^ rabbit polyclonal (1:1000; Millipore: ABN1381) and anti-pan TrkB mouse monoclonal (1:500; BioLegend: 695102) antibodies overnight at 4°C. Membranes were washed (4 washes, 10 min each) in TBS-Tween (TBST) and incubated with IRDye 800 CW goat anti-rabbit (1:15000; LI-COR Biosciences) and IRDye 680 RD goat anti-mouse (1:15000; LI-COR Biosciences) antibodies in blocking buffer (0.1% Tween 20; 0.1% SDS) for 1 hr. After incubation, membranes were washed in TBST followed by dH_2_O and were scanned on an Odyssey CLx multiplex fluorescence imaging system (LI-COR Biosciences). Blots were scanned at 700 and 800 nm simultaneously to allow direct comparisons of pan-TrkB and phospho-TrkB signals from the same sample, respectively.

After scanning for phospho- and pan-TrkB, the membranes were re-blocked and stained for tubulin. Briefly, membranes were reblocked for 1 hr in TBS Odyssey blocking buffer (LI-COR) and incubated with anti-β-tubulin mouse monoclonal antibody (1:30,000; #86298T, Cell Signaling) overnight at 4°C, followed by incubation with IRDye 680 RD goat anti-mouse antibody (1:15000; LI-COR 129 Biosciences) in blocking buffer (0.1% Tween 20; 0.1% SDS) for 1 hr. Membranes were washed in TBST, followed by dH_2_O, and scanned again on an Odyssey CLx fluorescence imaging system (LI-COR Biosciences) at 700 nm.

Several TrkB bands were noted on the blots (Supplementary Figure 1), most notably between 100-150kD and between 75-100 kD, suggesting relatively mature full-length (TrkB-FL) and truncated forms of TrkB (TrkB-T), respectively. We additionally observed bands between 25 and 37 kD, suggesting an intracellular cleavage product (e.g., Fonseca-Gomes et al., 2019). Bands corresponding to TrkB-FL were selected for quantification (see Supplemental Figure 1). Tubulin was used as a loading control for individual panTrkB and phosphoTrkB measurements.

Signals from western blots were quantified using Image Studio software (LI-COR Biosciences). Target bands were manually fitted and integrated optical density (OD) was measured with background correction using the median OD of pixels above and below each band. Only blots that had linear standard curves (R^2^ > .95) were included in the analysis (see Supplementary Figure 1).

### 2.9 Analysis and Statistics

#### 2.9.1. Maze training

Learning was quantified as the number of trials to reach a criterion of 9/10 correct trials with at least 6 consecutively correct in that span. Trial accuracy was defined as the number of correct choices in each block of 15 trials. Choice latency on the first trial was used to assess non-mnemonic effects of 7,8-DHF treatment related to maze training in general. Analyses of variance (ANOVAs) with Fisher’s LSD comparisons were used to determine significant differences among treatment groups (1, 5, and 10 µg 7,8-DHF and vehicle-treated conditions) in trials to criterion (TTC) and choice latency. A mixed model ANOVA was run to determine drug effects on choice accuracy across trial blocks; 7,8-DHF conditions were pooled together as trial accuracy was no different across all 7,8-DHF doses.

#### 2.9.2. TrkB activation

Values for pTrkB-FL and panTrkB-FL were normalized with tubulin as a loading control for each lane. To compare measures across blots and blotting conditions, values from experimental samples (30 µg loaded) were then normalized to the value from the pooled 30 µg standard included in each blot. As a measure of TrkB phosphorylation state, ratios of pTrkB-FL to panTrkB-FL ratios were determined for each individual lane.

Although total and phosphorylated levels of TrkB were analyzed independently, the primary outcome measure for TrkB activation state was pTrkB-FL relative to panTrkB-FL. One outlier for pTrkB/panTrkB was excluded based on the Grubb’s test; the values removed came from a 7,8-DHF-treated sample. Averages of values from sample duplicates were used for summary data. Independent samples t-tests were run to assess differences in panTrkb-FL, pTrkb-FL, and pTrkB/TrkB across treatment groups. All tests were run in SPSS (IBM) or Graphpad Prism with alpha set to 0.05.

## 3. Results

A single infusion of 7,8-DHF into the dorsolateral striatum 20 min before training robustly enhanced response learning. Compared to rats that received vehicle injections, intra-striatal infusions of each dose (1, 5, and 10 µg / hemisphere) of 7,8-DHF significantly reduced the number of trials needed to reach the learning criterion (F_3,12_ = 7.50, p < 0.05; each dose vs. vehicle-treated: p’s < 0.05; Figure 2A). Vehicle-infused rats took about twice as many trials (∼50 trials) to reach the learning criterion as did those infused with 7,8-DHF (range across doses: 23-30 trials). There were no significant differences in trials to criterion across 7,8-DHF doses (p’s > 0.1).

**Figure 2.**
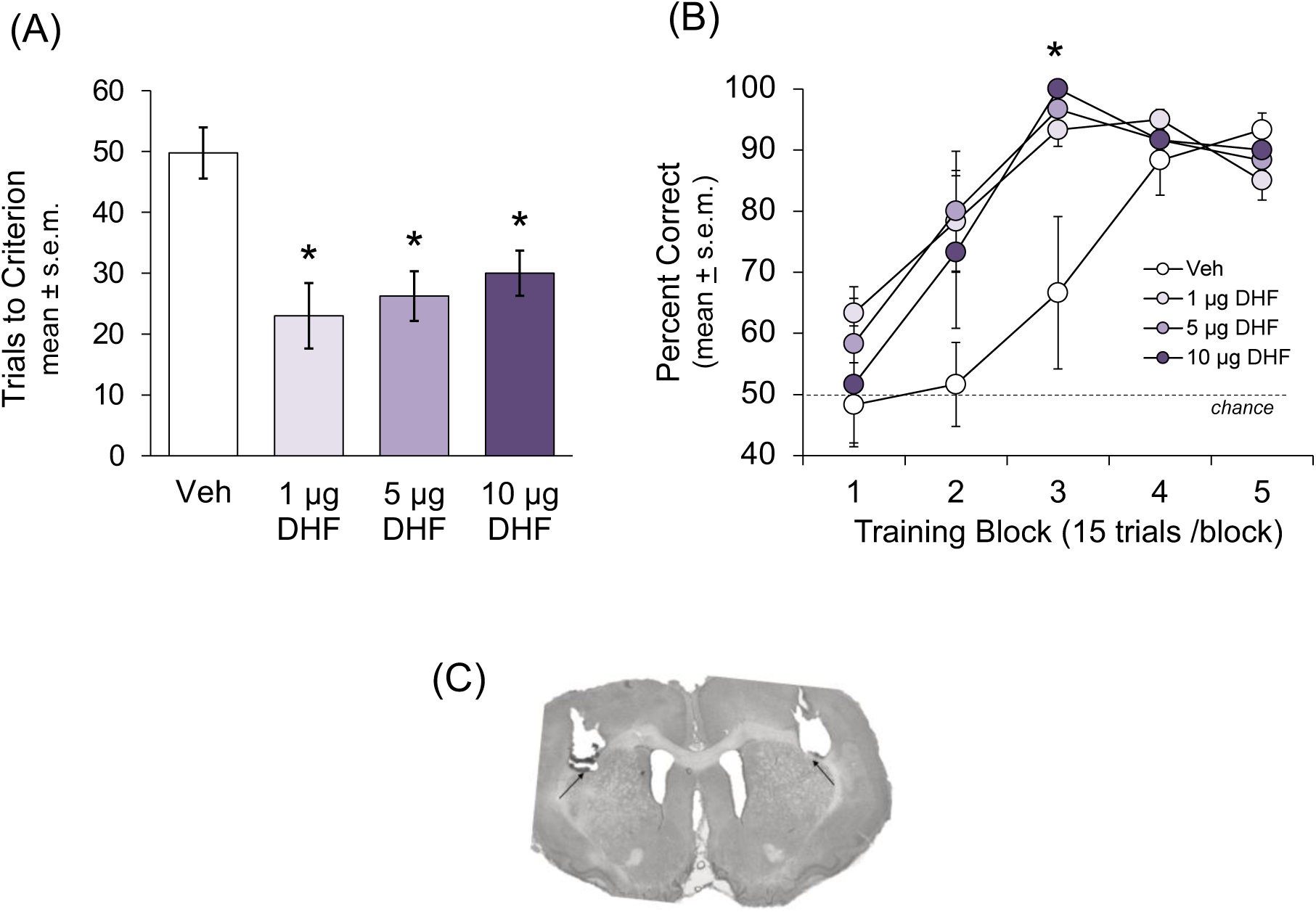
Effects of 7,8-DHF infusions into the dorsolateral striatum on response learning. The three doses of 7,8-DHF tested (1, 5, or 10 µg/hemisphere) each enhanced response learning as compared to that in vehicle-controls, as shown by 7,8-DHF-induced reduction in *(A)* TTC, and *(B)* trial accuracy across the training session (5 blocks of 15 trials/each). The dashed line indicates chance levels of choice accuracy (50%). *p < 0.05 vs DMSO vehicle. n = 4 in each group. *(C)* Micrograph showing correct placements of infusion cannulae in the dorsal striatum. Arrows point to tip of injection site.

On measures of trial accuracy across training, rats that received 7,8-DHF learned more quickly than did vehicle-treated rats. While rats in all groups started the task at or near chance levels (50% trial accuracy), rats in the 7,8-DHF groups reached > 75% accuracy by trial block 2 and > 95% accuracy by trial block 3, compared to 52% and 67% in vehicle-treated rats, respectively. Consistent with these observations, ANOVA tests on trial accuracy (treatment group x training blocks) revealed significant main effects of treatment (F_1,14_ = 9.68, p < 0.01) and training block (F_4,56_ = 26.07, p < 0.01) as well as a significant interaction between treatment and training block (F_4,56_ = 5.122, p < 0.01; Figure 2B). There were no significant differences (p’s > 0.10) between the three 7,8-DHF doses in choice accuracy. Choice latencies during the first trial of training ranged on average from 26 to 41 sec and were comparable across all conditions (F_3,12_ = 0.44; p > 0.70; data not shown).

Representative cannula placements are shown in Figure 2C. Except for two rats (1 = VEH, 1 = 1 µg 7,8-DHF), which were excluded from further analysis, all injection sites were within the dorsolateral striatum.

Protein levels were assessed in striatal samples collected from rats 20 min after 7,8-DHF infusions, a time matching when training began in rats tested behaviorally.

There was a significant ∼43% increase in the ratio of pTrkB-FL to panTrkB-FL in 7,8- DHF-treated rats compared to vehicle-treated controls (t_13_ = 2.49; p < 0.02; Figure 3A, B), suggesting that 7,8-DHF infusions into the dorsolateral striatum activated TrkB. Levels of panTrkB-FL were approximately 20% lower in DHF-treated rats than in controls (p < 0.06) and pTrB-FL levels were about 20% higher in 7,8-DHF-treated rats than in controls (p < 0.2), although these effects did not reach statistical significance (Figure 3C, D). Tubulin levels were not themselves significantly altered by 7,8-DHF treatment (p’s > 0.05; data not included).

**Figure 3.**
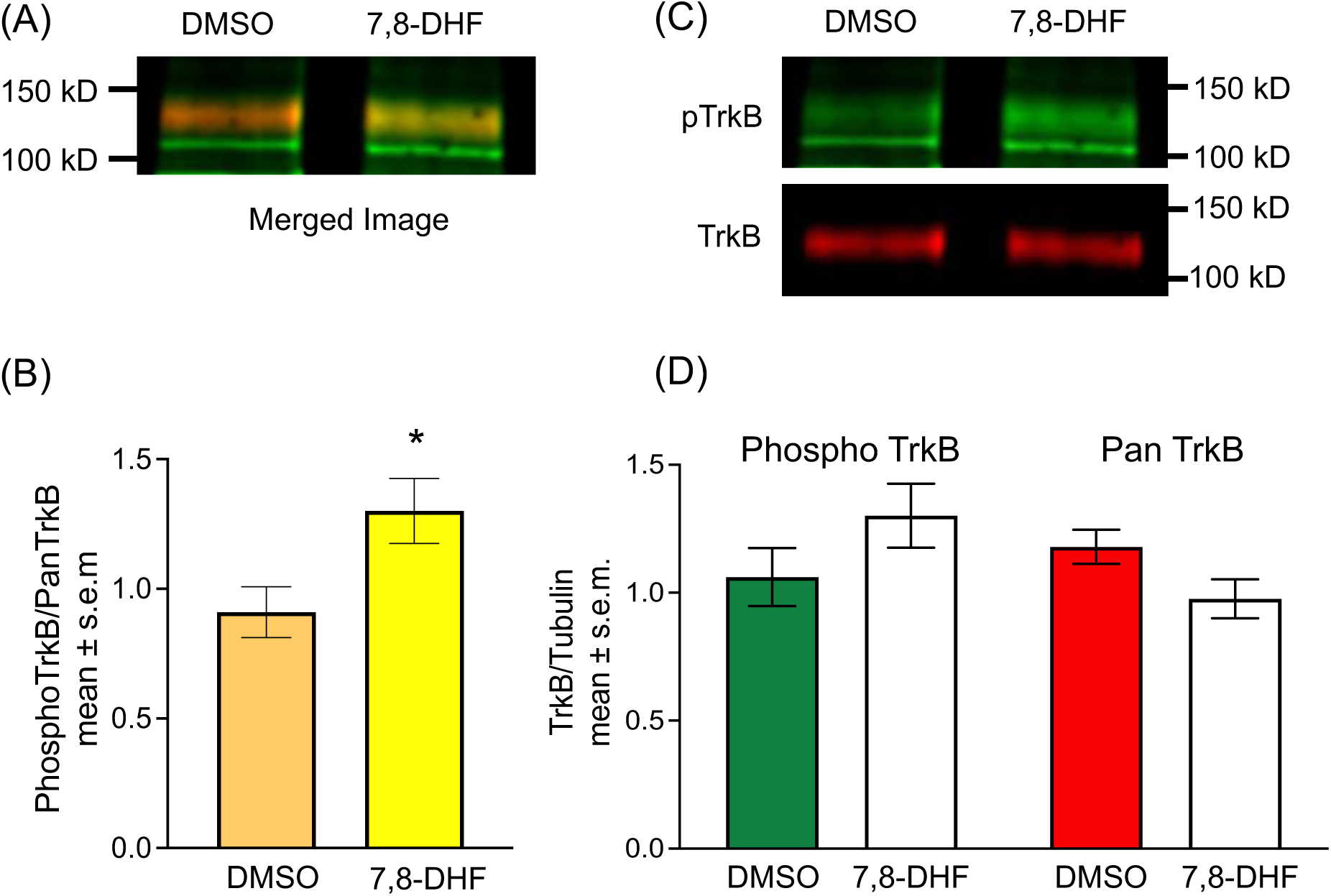
*Left panel. (A)* Merged image of 100-150 kD bands following 700 nm and 800 nm scans. Yellow indicates areas of overlap between phosphorylated and total TrkB. *(B)* The ratio of phosphorylated t TrkB^816^- to total TrkB-FL in the dorsal striatum increased significantly after 7,8-DHF infusions, suggesting heightened activation of TrkB. *p < 0.05 vs. DMSO vehicle. n = 8 in each group. *Right panel. (C)* Representative bands showing increased pTrkB^816^ (green) relative to panTrkB (red) in DHF-treated rats. *(D)* Graph of total and phosphorylated TrkB^816^ in the dorsolateral striatum after local infusions of 7,8-DHFrelative to tubulin loading control. Though not statistically significant, 7,8-DHF infusions decreased total TrkB and increased phosphorylated TrkB^816^ compared to DMSO infusions.

## 4. Discussion

Infusions of the selective TrkB receptor agonist 7,8-DHF into the dorsolateral striatum significantly and robustly accelerated learning on the response task. Enhancement of learning was seen at all tested doses, which spanned a log unit. Importantly, all groups performed at chance levels and displayed similar trial latencies at the start of training, suggesting that 7,8-DHF enhancement of learning measures was likely due to changes in learning and memory processes rather than non-mnemonic factors such as locomotor or exploratory behaviors. In the DHF groups, enhanced learning was evident as early as the second trial block (trials 16-30). By Block 4, the vehicle-treated group expressed good learning, approaching the percent correct in the 7,8-DHF-treated groups. Consistent with results obtained before (Andero et al., 2011; Bollen et al., 2013; Rawlings-Mortimer et al., 2023), the enhancement of learning in the present experiment was evident in young adult rats without cognitive impairment. The learning scores of the rats tested here with implanted cannulae were comparable to those we have seen previously in unimplanted young adult rats (e.g., Gardner et al., 2022). Together with the wealth of evidence that TrkB mediates hippocampal learning and memory (for reviews see: Minichiello, 2009; Yamada and Nabeshima, 2003), the findings reported here support a role for TrkB-dependent mechanisms in mnemonic processes across multiple memory systems, specifically extending the role of TrkB mechanisms to cognitive processing in the striatum.

Enhancement of response learning by 7,8-DHF was accompanied by TrkB activation in the striatum. As reported before in a chronic treatment model (Zeng et al., 2012a), 7,8-DHF induced rapid effects on TrkB tyrosine 515 and 816 phosphorylation. These findings suggest a key role for striatal TrkB activation in the augmentation of striatum-sensitive learning processes and extend prior reports showing a correlation between activated TrkB and response learning efficacy (Pahng and Colombo, 2017). TrkB phosphorylation after treatment with 7,8-DHF results in downstream activation of cAMP responsive element binding protein (CREB) (Agrawal et al., 2015; Ahmed et al., 2021; Yang et al., 2014), particularly after TrkB^816^ activation (Minichiello et al., 2002). Phosphorylation of CREB is an important component of the molecular events associated with memory and neuronal plasticity (for reviews, see: Kaushik et al., 2022; Lisman et al., 2018; Silva et al., 1998; Yin and Tully, 1996), including CREB involvement in learning processes within the striatum (Brightwell et al., 2008; Colombo et al., 2003; Kathirvelu and Colombo, 2013; Qian et al., 2015). Of interest, several treatments that enhance cognition, including glucose and lactate (Griego and Galvan, 2023; Hertz and Chen, 2018; Hossain et al., 2020), estrogens (Harte-Hargrove et al., 2013), glucocorticoids (Chen et al., 2017; Finsterwald and Alberini, 2014; Jeanneteau et al., 2019; Notarus and van den Buuse, 2020), and physical and cognitive exercise (Griesbach et al., 2009; Korol et al., 2013) each appear to enhance memory by activation of BDNF, TrkB receptors, and CREB.

As 7,8-DHF induced TrkB-FL activation 20 min after the infusions, corresponding to the onset of training, our findings additionally suggest that signaling through TrkB at the time of training confers enhancements in learning abilities. Most studies of 7,8-DHF enhancement of cognitive functions have use prolonged administration of the drug, generally for many days or weeks (Agrawal et al., 2015; Bryant et al., 2021; Castello et al., 2014; Chen et al., 2023; Dhaliwal et al., 2024; Seese et al., 2020; Schultz and Crawley, 2020; Tian et al., 2015; Zeng et al., 2012a,b). Our results showing that a single drug treatment given near the time of training effectively enhanced learning are consistent with the findings of Andero et al. (2011) and Bollen et al. (2013). The present findings are also consistent with prior work showing that inhibition of hippocampal or striatal TrkB activation at the time of place or response training blocked learning enhancements observed after priming experiences, i.e., a 20-min test on a working memory task or three weeks use of running wheels (Korol et al., 2013). Collectively, these results suggest that enhancement of learning and memory may be mediated by acute release of BDNF and acute activation of TrkB receptors. Thus, while trophic mechanisms such as neurogenesis may also be important contributors to BDNF functions (e.g. Bekinschtein et al., 2011; Kim and Park, 2018; Liu and Nusslock, 2018; Vivar et al., 2013), the acute effects are sufficient to mediate BDNF enhancement of learning and memory in the absence of these longer-term changes. A hypothesis that TrkB activation mediates rapid effects on cell-cell signaling contributing to learning and memory is further supported by several findings. TrkB activation rapidly increases Ca^2+^ mediated hippocampal synaptic plasticity by regulating membrane excitability (Hammond et al., 2006) and postsynaptic potentials on a time scale consistent with glutamatergic signaling (Kafitz et al., 1999); indeed, the modulatory role of BDNF/TrkB signaling in postsynaptic transmission is explained in part by relatively rapid effects on excitatory NMDA glutamatergic receptors and inhibitory GABA receptors altering synaptic excitatory tone (Rose et al., 2004).

Thus, the findings of the present study extend 7,8-DHF and, by extrapolation, TrkB activation by BDNF enhancement of learning and memory beyond the hippocampus to the striatum. In addition, these effects are seen with an acute treatment near the time of training, supporting the view that training-related release of BDNF may mediate enhancement of cognitive processes in the striatum. Moreover, the results here show that 7,8-DHF can enhance learning and memory in rats with intact abilities. With the extensive evidence that BDNF mediates the effects of exercise on learning and memory (Cefis et al., 2023; Jaberi and Fahnstock, 2023), the present results suggest that 7,8-DHF, as a mimic of BDNF, may be useful as an alternative to exercise for the augmentation of multiple cognitive functions. This may prove particularly important for populations with limited access to physical activity.

## CRediT authorship contribution statement

**Robert S. Gardner**: Investigation, Formal analysis, Writing - review & editing, Visualization. **Matthew T. Ambalavanar**: Investigation, Formal analysis, Writing - review & editing, Visualization. **P.E. Gold**: Conceptualization, Formal analysis, Writing - original draft, Writing - review & editing, Visualization, Supervision, Project administration, Funding acquisition. **D.L. Korol**: Conceptualization, Formal analysis, Writing - original draft, Writing - review & editing, Visualization, Supervision, Project administration, Funding acquisition.

## Declaration of Competing Interest

The authors declare that they have no known competing financial interests or personal relationships that could have appeared to influence the work reported in this paper.

## Acknowledgements

This work was supported by NSF IOS 13-18490, NIDA DA038798, NIA AG057947, and by P30 AG034464 through the Center on Aging and Policy Studies at Syracuse University.

---

**Supplementary Figure 1.**
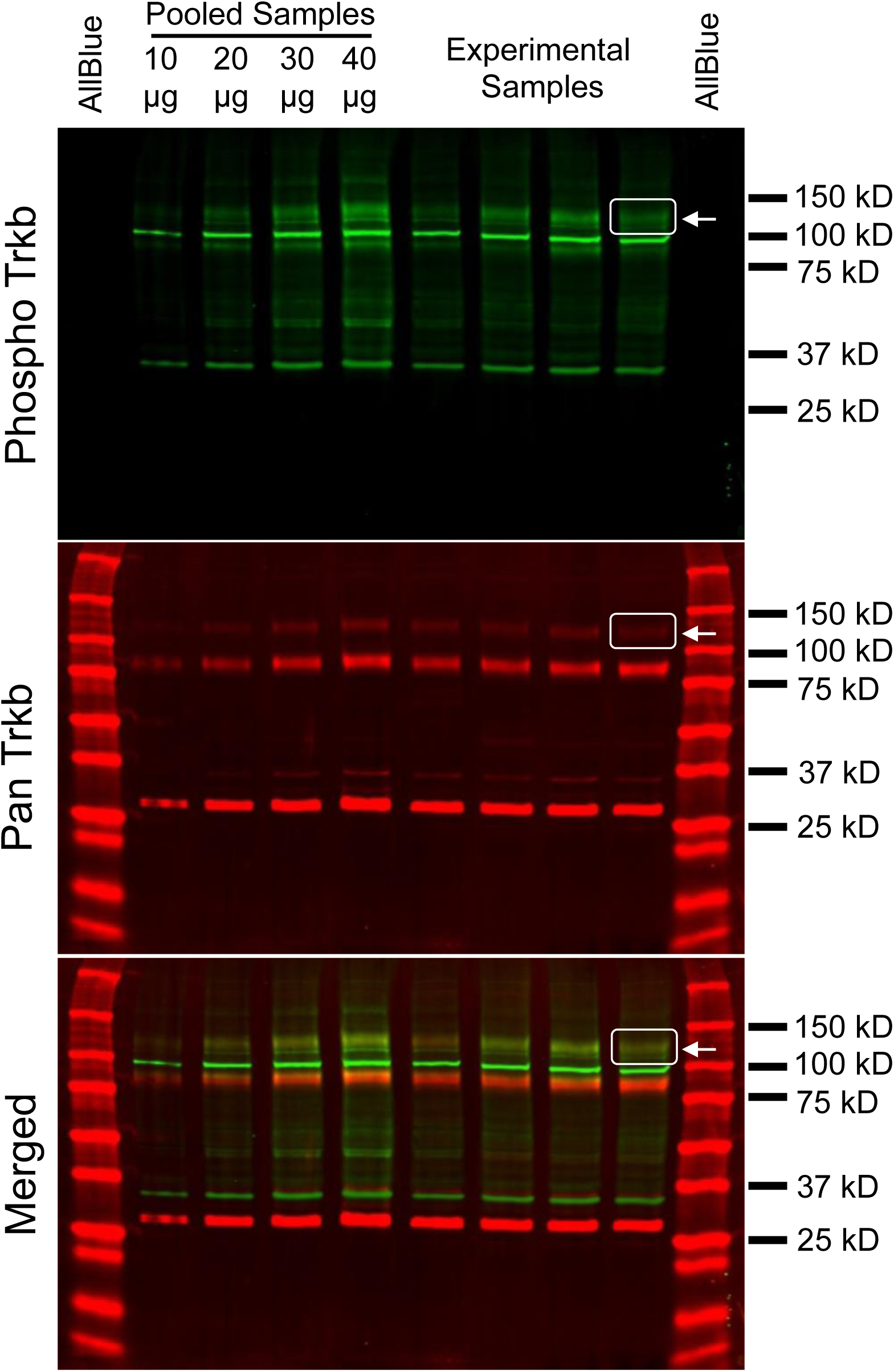
Representative blots for pTrkB^816^ (green) and panTrkB (red) following scanning with near infrared detection at 700 nm and 800 nm (Odyssey CLx, LICOR). All blots included pooled standard samples spanning 10 µg, 20 µg, 30 µg, and 40 µg loaded protein, four experimental samples (30 µg loaded protein), and two MW ladders. Several bands were detected, most notably those between 100-150kD and between 75-100 kD, believed to represent mature full-length (TrkB-FL) and truncated (TrkB-T) forms of TrkB, respectively. Another set of bands was found between 25 and 37 kD, which may reflect an intracellular cleavage product (Fonseca-Gomes et al., 2019). Bands identified by white arrows and boxes were those included for analysis. Note the linearity in the pTrkB-FL and panTrkB-FL signals from the standards, a property that was used for inclusion in data analysis. All blots were linear (R^2^ > 0.95) across 10-40 µg loaded protein except for tubulin measures on two blots, which were both linear (R^2^ > 0.98) across 10-30 µg.

## Notes

### Competing Interest Statement

The authors have declared no competing interest.

